# Distinct networks coupled with parietal cortex for spatial representations inside and outside the visual field

**DOI:** 10.1101/2020.07.22.215517

**Authors:** Bo Zhang, Fan Wang, Qi Zhang, Naya Yuji

**Author notes:** **Corresponding author:** Yuji Naya, Address: School of Psychological and Cognitive Sciences, Peking University, No. 52, Haidian Road, Wang Kezhen Building, Room 1707, Haidian District, Beijing 100805, China.

## Abstract

Our mental representation of egocentric space is influenced by the disproportionate sensory perception of the body. Previous studies have focused on the neural architecture for egocentric representations within the visual field. However, the space representation underlying the body is still unclear. To address this problem, we applied both fMRI and MEG to a spatial-memory paradigm by using a virtual environment in which human participants remembered a target location left, right, or back relative to their own body. Both experiments showed larger involvement of the frontoparietal network in representing a retrieved target on the left/right side than on the back. Conversely, the medial temporal lobe (MTL)-parietal network was more involved in retrieving a target behind the participants. The MEG data showed preferential connectivity in the alpha-band frequency in both networks. These findings suggest that the parietal cortex may represent the entire space around the self-body by coordinating two distinct brain networks.

## Introduction

While planning to reach a target, it is necessary to obtain its location information in the body-centered reference frame or the so-called “egocentric” spatial coordinates^1–4^. Previous studies using humans and nonhuman primates indicated the crucial involvement of the parietal cortex in the representation of egocentric location for sensory perception, motor action, and their coordination^4–10^. Anatomically, the parietal cortex is located at the late stage of the dorsal pathway, which is often named the “where” or “how” pathway^11–12^. Perceptual signals, including visual, somatosensory, and vestibular information, converge on the parietal cortex, which interacts with the frontal eye field (FEF), the region associated with attention and eye movement toward a target in the external space around the self-body^13–16^. These anatomical connection patterns are consistent with those reported by neuropsychological studies, that is, damage to the parietal cortex results in a form of impaired egocentric spatial awareness known as hemispherical neglect, which leads to neglect of a target on one side of the field of vision^17^. Hemispherical neglect appears not only in perception but also in the memory field, where it is known as “representational neglect”^18^. On the basis of data accumulated from human imaging studies examining spatial navigation and episodic recollection^19–23^, the parietal cortex is considered to represent the egocentric space for both perception and memory.

Mnemonic representations of the egocentric space raise the question of whether the spatial representation differs between the inside and outside of a visual field. While the former can be represented in both vision and memory, the latter can be represented only in memory. Previous human behavioral studies reported decreased performance^24^ or prolonged response latency^25^ when participants located a target behind them. This phenomenon, known as the “alignment effect”^26^ or “front facilitation,”^27^ suggests that the space surrounding our body is coded heterogeneously by different neural systems depending on the target location along the anterior-posterior axis of the self-body^27^. To explore the neural architectures responsible for the representation of egocentric space, previous studies mostly examined the target representation within the visual field; however, one neuroimaging study, which examined target representation retrieved from memory, suggested that the parietal cortex codes the whole space around the self-body^28^. However, the representation properties of the inside/outside of the visual field in the parietal cortex and its interactions with associated brain areas remain unsolved.

To characterize the neural architectures supporting the mental representations of egocentric space inside and outside the visual field, we applied functional magnetic resonance imaging (fMRI) and magnetoencephalography (MEG) to a spatial-memory paradigm using a 3D virtual environment for human participants (Fig. 1a), which we recently devised^29^. In this paradigm, participants encoded a spatial relationship among three objects (walking period) in each trial. While the same three objects were used across trials, their spatial relationships differed pseudorandomly. After the walking period, one of the three objects was presented in the center of the display with the environmental background, which prompted the participants to feel as if they were facing the objects in the virtual environment (facing period)^29^. Then, one of the other two objects was presented on a scrambled background as the target object (targeting period). The participants remembered the location of the target object relative to the body. The to-be-remembered targeting locations could be in the left, right, or back positions of the participants’ egocentric spaces (Fig. 1b). This spatial-memory paradigm allowed us to compare the neural representations of egocentric space between the inside and outside of the visual field when the participants remembered a target location, which had been encoded in each trial.

**Figure 1.**
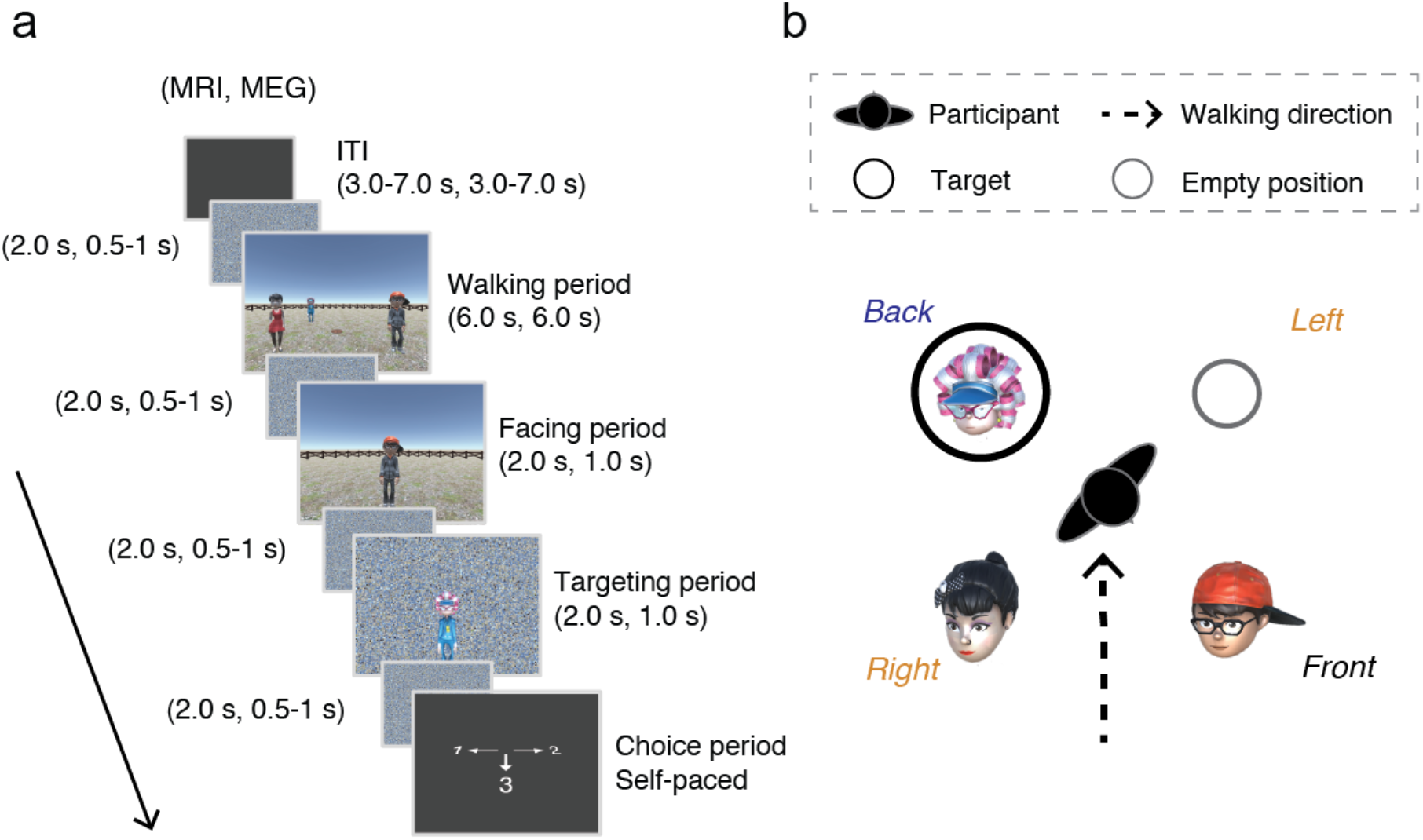
a) Spatial-memory task paradigm. Each trial consisted of four periods. Walking period: participants walked toward three human characters using the first-person perspective and stopped on a wood plate in the center. Facing period: one of the human characters was presented, indicating the participant’s current self-orientation. Targeting period: a photo of the target character was presented on a scrambled background. Choice period: the participants chose the direction of the target character relative to their body upon the presentation of a response cue. b) The spatial relationship between the participant and the human characters in the example trial of Fig. 1a, with the target location behind the participant’s self-body (*black circle*).

## Results

### Behavioral performance

Nineteen and 12 healthy volunteers participated in the fMRI and MEG experiments, respectively. The participants performed the spatial-memory task with high accuracy rates in both the experiments (MRI: 93.6% ± 1.5%; MEG: 90.4% ± 1.9%). The performances did not differ among the three target locations (i.e., right, left, and back) in both fMRI [*F* (2,54) = 0.82, *P* = 0.44] and MEG [*F* (2,33) = 0.08, *P* = 0.93] experiments (Fig. 2a), indicating that the participants solved the spatial memory task accurately regardless of the target location.

**Figure 2.**
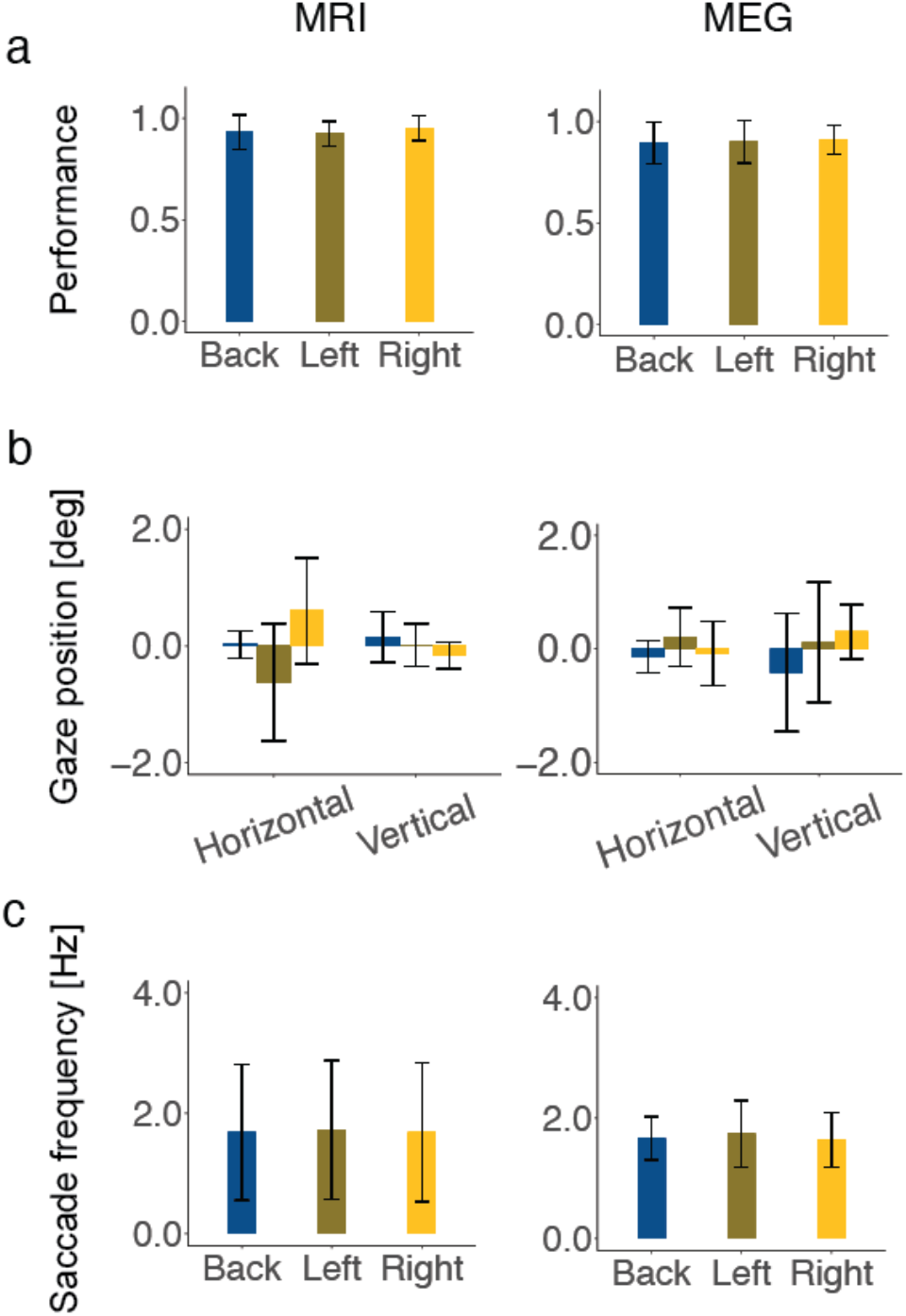
a) Performance of participants for the three egocentric target locations in the MRI (n = 19) and MEG (n = 12) experiments. A statistically significant difference was not detected in the performances of the target locations by repeated-measures one-way ANOVA in the MRI [*F*(2,54) = 0.82, *P* = 0.44] or MEG [*F*(2,33) = 0.08, *P* = 0.93] experiments. b) Gaze positions of the participants during the targeting period (4 s for MRI, 1 s for MEG) were shown for each target condition. No significant difference was found among the left, right, and back-target conditions in the MRI experiments [horizontal: *F*(2, 51) = 1.7, *P* = 0.19, vertical: *F*(2, 51) = 0.08, P = 0.92) or the MEG experiments [horizontal: *F*(2, 27) = 0.32, *P* = 0.73, vertical: *F*(2, 27) = 0.42, *P* = 0.66]. c) No significant difference in the frequencies of saccadic eye movements was found among the three conditions during the targeting period [MRI: *F*(2, 51) = 0.01, *P* = 0.99; MEG: *F*(2, 27) = 0.14, *P* = 0.87]. Error bars indicate standard deviations.

### MRI contrast analysis

The neural activity in the fMRI experiment was examined using the 4-s time-window of the targeting period in which the participants remembered the location of the target object (i.e., human character) relative to their self-body in the virtual environment. A contrast analysis showed a significantly stronger blood-oxygen-level-dependent (BOLD) signal for the left/right-target condition than for the back-target condition in the parietal cortex, precuneus, FEF, and SMA (Fig. 3a, *P* < 0.01; initial threshold, *P* < 0.05; cluster-corrected for multiple comparisons). In contrast, no brain areas showed a significantly stronger BOLD signal for the back-target condition than for the left/right-target condition in the same statistical condition with multiple comparisons, although we found a significant cluster for the back-target location in the right rhinal cortex (entorhinal cortex [ERC] and perirhinal cortex [PRC]) of the MTL when we used a more liberal threshold (Fig. 3b, *P* < 0.05, uncorrected).

**Figure 3.**
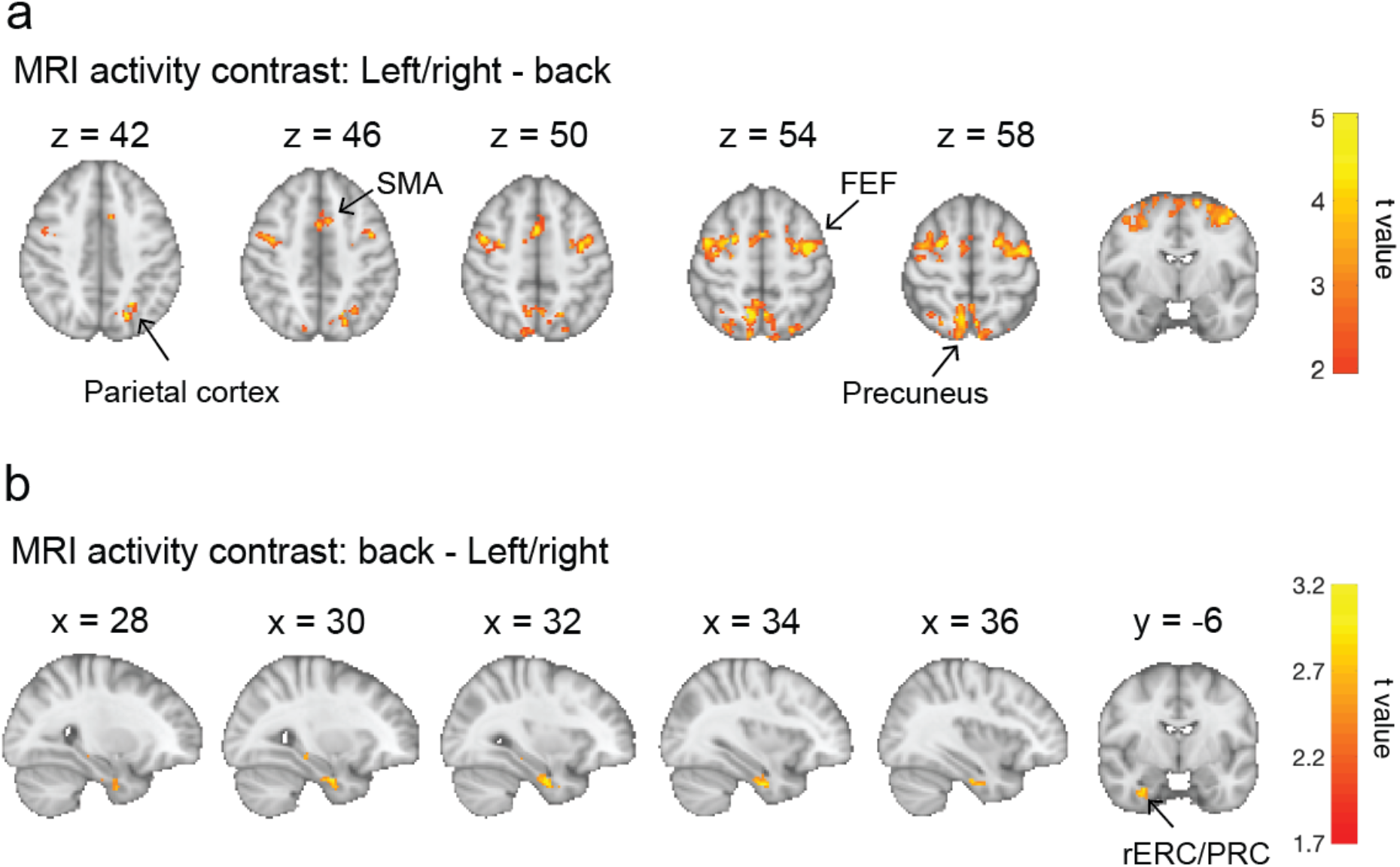
a) MRI contrast for the “left/right-target” – “back-target” condition: significant clusters were observed in the parietal cortex, precuneus, FEF, and SMA (*P* < 0.01, initial threshold, *P* < 0.05, cluster-corrected for multiple comparison). b) MRI contrast for the “back-target” – “left/right-target” condition: a cluster was observed in the right rhinal cortex, including the entorhinal cortex (ERC) and perirhinal cortex (PRC) (*P* < 0.05 uncorrected).

Regarding the larger activation of the dorsal brain regions under the left/right-target conditions than the back-target condition, one question might be whether the elevated activities can be explained by eye movements under different target conditions. To test this possibility, we examined the participants’ gaze positions during the 4-s time window of the targeting period. The gaze positions did not differ among the egocentric locations of the retrieved targets in the horizontal direction [*F* (2, 51) = 1.7, *P* = 0.19] or the vertical direction [*F* (2, 51) = 0.084, *P* = 0.92] (Fig. 2b). In addition to the average gaze position, we examined the frequencies of saccadic eye movements (see Methods), but no significant difference was found between the left/right and back-target conditions during the targeting period [*F* (2, 51) = 0.007, *P* = 0.99] (Fig. 2c). These results indicate that the increased activity in the dorsal brain regions was not due to eye movements that may have been induced by the retrieved egocentric locations. Taken together, these findings reasonably suggest that the frontoparietal cortical areas were involved in an additional mental process to represent a retrieved target to the left/right in comparison with that behind the participants’ self-body. Similarly, remembering a target behind the self-body may employ the right MTL more than when the participants remembered a target to their left/right.

### MRI functional-connectivity analysis

We subsequently investigated the brain regions that interacted with the parietal cortex and precuneus when the participants remembered the target location to their left, right, and back. To examine the interactions, we conducted whole-brain functional connectivity analysis using the anatomical ROIs of the parietal cortex and precuneus as the seeds based on the AAL (see Methods for details). The results showed that, in addition to a mutual connectivity between these brain areas, both the parietal cortex and precuneus exhibited a significantly higher connectivity to the FEF and SMA under the left/right-target condition than under the back-target condition (Fig. 4a, *P* < 0.01, initial threshold, *P* < 0.05, cluster-corrected for multiple comparison). This result indicates the cooperative involvement of the FEF, SMA, parietal cortex, and precuneus while processing a target object in the visual field, even when the egocentric object location was not perceived. In contrast, the parietal cortex, but not the precuneus, showed increased functional connectivity to the right ERC when a retrieved target location was behind the participants’ body (Fig. 4b, *P* < 0.01, initial threshold, *P* < 0.05, cluster-corrected for multiple comparisons via small-volume correction using the bilateral MTL mask). We also examined the functional connectivity using FEF and SMA seeds and found that these two brain regions exhibited strong connectivity to the parietal cortex and precuneus in the left/right-target condition but did not show increased functional connectivity to any other brain regions, including the ERC, in the back-target condition. These results suggest that our egocentric space is represented by two distinct brain networks: the frontoparietal network for the target within the visual field, and the MTL-parietal network for the target outside the visual field.

**Figure 4.**
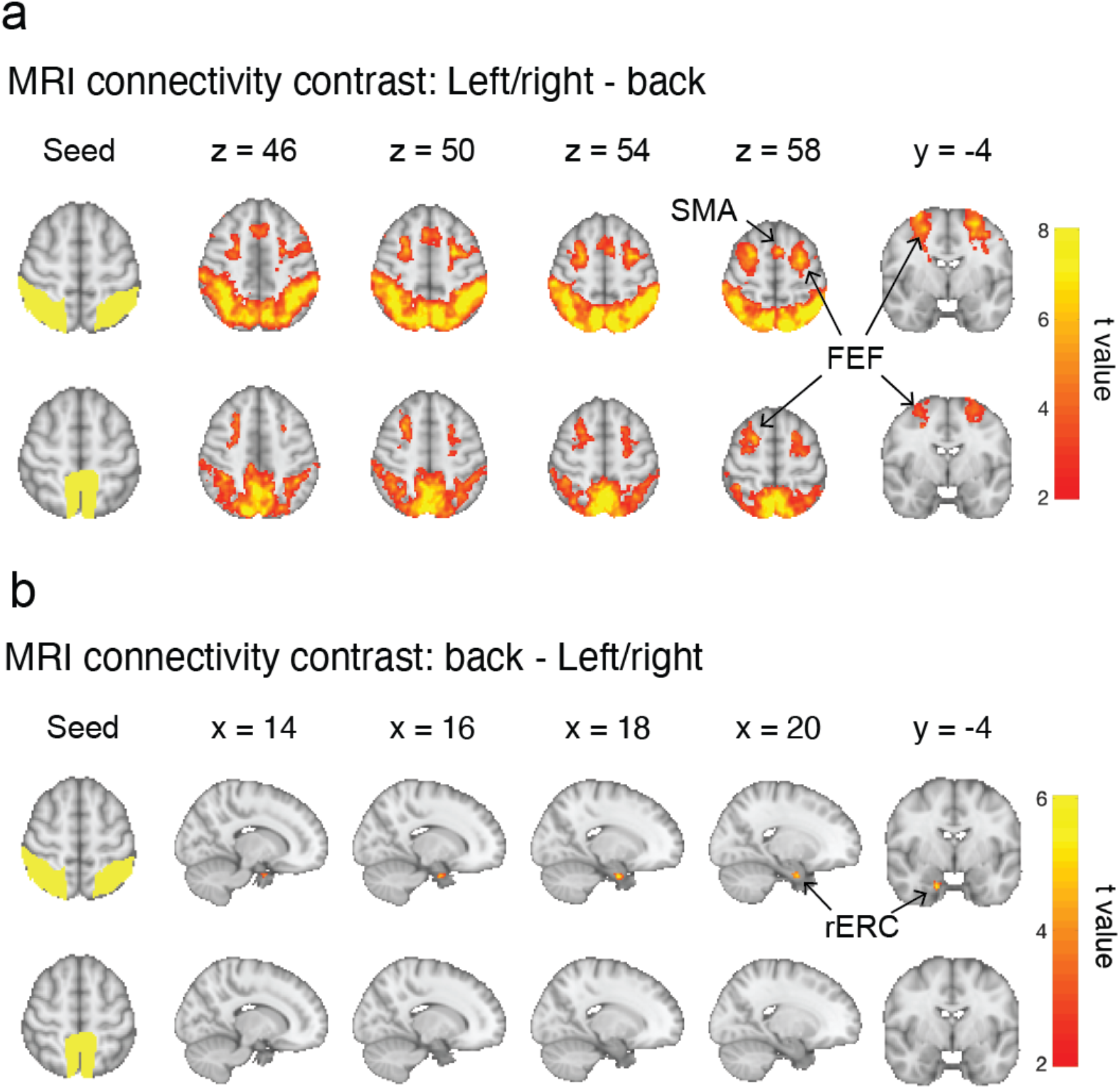
MRI connectivity analysis using the parietal cortex and precuneus as seeds. a) Connectivity contrast for the “left/right-target” – “back-target” condition: significant connections were found in the parietal cortex, precuneus, FEF, and SMA for both seeds (*P* < 0.01, initial threshold, *P* < 0.05, cluster-corrected for multiple comparison). b) Connectivity contrast for the “back-target” – “left/right-target” condition: a significant connection was observed between the parietal cortex and right ERC (*P* < 0.01, initial threshold, *P* < 0.05, cluster-corrected for multiple comparisons via small-volume correction using the bilateral MTL mask); no connection was found between the precuneus and MTL regions.

### MEG contrast analysis

The fMRI experiments showed preferential involvement of the frontoparietal network in the left/right-target condition and those of the MTL-parietal network in the back-target condition. To examine the temporal dynamics of these brain networks, we subsequently conducted an MEG study using the same spatial memory task that was used for the fMRI study, except for the time parameters (Fig. 1a).

Figure 5a shows the results of the contrast analysis that compared the activity strength at each sensor between the left/right-target and back-target conditions. We found a cluster of sensors in the left-posterior area, which showed significantly stronger activity under the left/right-target condition than under the back-target condition during 0.67–0.85 s after the onset of the targeting period (*P* < 0.05, initial threshold, two tailed; *P* = 0.04, spatial-temporal cluster-corrected for multiple comparison) (Fig. 5b). Conversely, no cluster of sensors showed stronger activity under the back-target condition than under the left/right-target condition. In addition to the fMRI study, we examined the participants’ gaze positions and frequencies of saccadic eye movements in MEG experiments. There was no significant difference among the left, right, and back target conditions during the 1-s time-window of the targeting period in the eye positions [*F* (2, 27) = 0.32, 0.42, *P* = 0.73 and 0.66, respectively, for the horizontal and vertical positions, respectively] (Fig. 2b) nor the saccade frequencies [*F* (2, 27) = 0.14, *P* = 0.87] (Fig. 2c). These results were consistent with the results of the fMRI experiments, indicating predominant spatial representation processing within the visual field relative to that outside the visual field.

**Figure 5.**
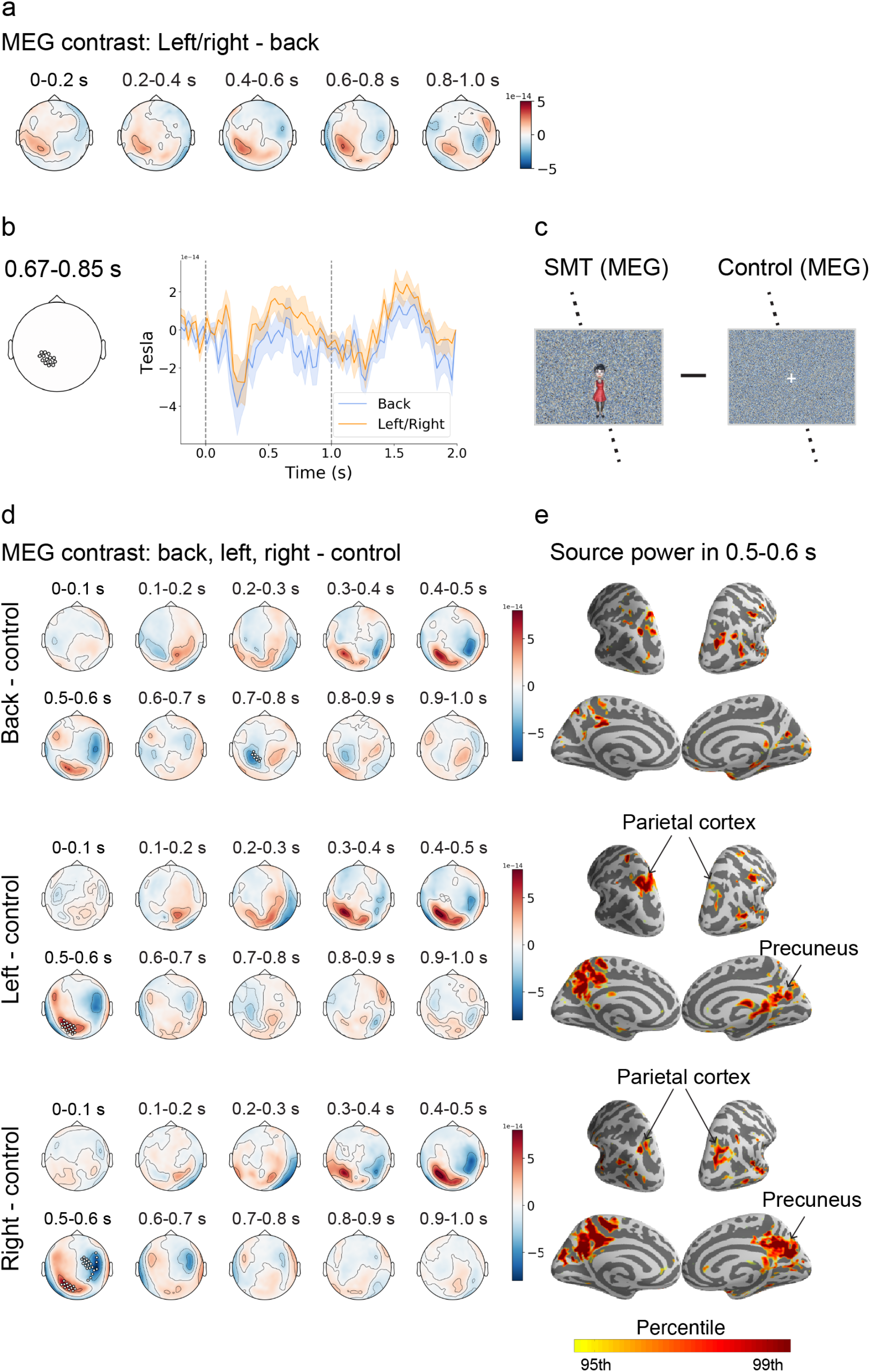
a) Mean topographic map for the MEG contrast of “left/right-target” – “back-target” condition for every 0.2 s during the targeting period. b) Time courses of the signal strength on the left-posterior cluster of sensors in the back-target and left/right-target conditions. A significantly higher activity was found for the left/right-target condition relative to the back-target condition during 0.67–0.85 s after the onset of the targeting period (*P* < 0.05, initial threshold, *P* < 0.05, spatial-temporal cluster correction for multiple comparison, two tailed). c) A comparison between the spatial-memory-task (SMT) trials and the control-condition trials in the MEG experiment. d) Mean topographic map for the left-target, right-target, and back-target conditions relative to the control condition for every 0.1 s during the targeting period. Significant clusters were found in the left-posterior from 0.5 to 0.6 s for the left-target and right-target conditions (*P* < 0.05, initial threshold; *P* < 0.05, cluster-corrected for multiple comparison, two tailed), but not for the back-target condition. e) Source-power distribution on the brain surface for each of the three conditions relative to the control condition within 0.5–0.6 s. Color bar represents the percentile rank of source-power strength.

To examine the changes in neural activity required for the retrieval of a target location, we compared the activity strength under back-target, left-target, and right-target conditions with the control trials, in which a targeting-character was not presented and the participants were not required to remember a target location (Fig. 5c). Figure 5d shows the time course of the topographic activity map under each condition relative to that under control conditions. Increased activity was observed in the left posterior area of the head. This left-posterior cluster showed a significant increase in activity during 0.5–0.6 s from the onset of the targeting period under the left-target and right-target conditions (*P* < 0.05, initial threshold; *P* < 0.05, cluster-corrected for multiple comparison) although the same trend of increase in activity was observed under the back-target condition [*t*(11) = 2.42, *P* = 0.03, uncorrected]. To localize the brain regions contributing to the significant activity increase in the left-posterior cluster, we conducted a source analysis of the MEG signal^30^. The source powers were distributed largely in the parietal cortex and precuneus in the left- and right-target conditions (Fig. 5e). We also found that the source power for the back-target condition was distributed in the parietal cortex and precuneus, although the level of source power was smaller than those under the left -and right-target conditions.

To explore the brain regions exhibiting larger neural activity under the back-target condition than under the left/right-target condition, we conducted a whole-brain analysis to compare the source power between the two conditions at intervals of 0.2 s after the onset of the targeting period (Fig. 6a). We found a strong source-power for the back-target condition in the right MTL, including the ERC, during 0.2–0.4 s after the onset of the targeting period. Using anatomical ROIs of each hemisphere of the whole MTL, we examined the precise time courses of the source power for the back -and left/right-target conditions relative to the control trials. The results indicated elevations in the source power after the onset of the targeting period under both back-target and left/right-target conditions in both hemispheres of the MTL. However, only the right MTL exhibited a significantly larger source power for the back-target condition than for the left/right-target condition in the early phase (0.25–0.37 s) after the onset of the targeting period (Fig. 6b, *P* < 0.05, initial threshold; *P* < 0.05, spatial-temporal cluster correction for multiple comparison, two-tailed). We further examined the source power in the right MTL by using the anatomical masks of its subregions and found that the source power was larger under the back-target condition than under the left/right-target condition in all the subregions [HPC: *t*(11) = 3.00, *P* = 0.048; PHC: *t*(11) = 2.98, *P* = 0.049; PRC: *t*(11) = 3.22, *P* = 0.032; ERC: *t*(11) = 3.39, *P* = 0.024, Bonferroni-corrected for multiple comparisons (n = 4)] (Fig. 6c). Collectively, the MEG contrast analyses in source-power between the back- and left/right-target conditions indicate that the right MTL, including the ERC, was involved more in the back-target condition than in the left/right-target condition in the early phase (0.25–0.37 s) after the onset of the targeting period, while the parietal cortex and precuneus were involved more in the left/right-target condition than in the back-target condition in the late phase (Fig. 5b, 0.67–0.85 s).

**Figure 6.**
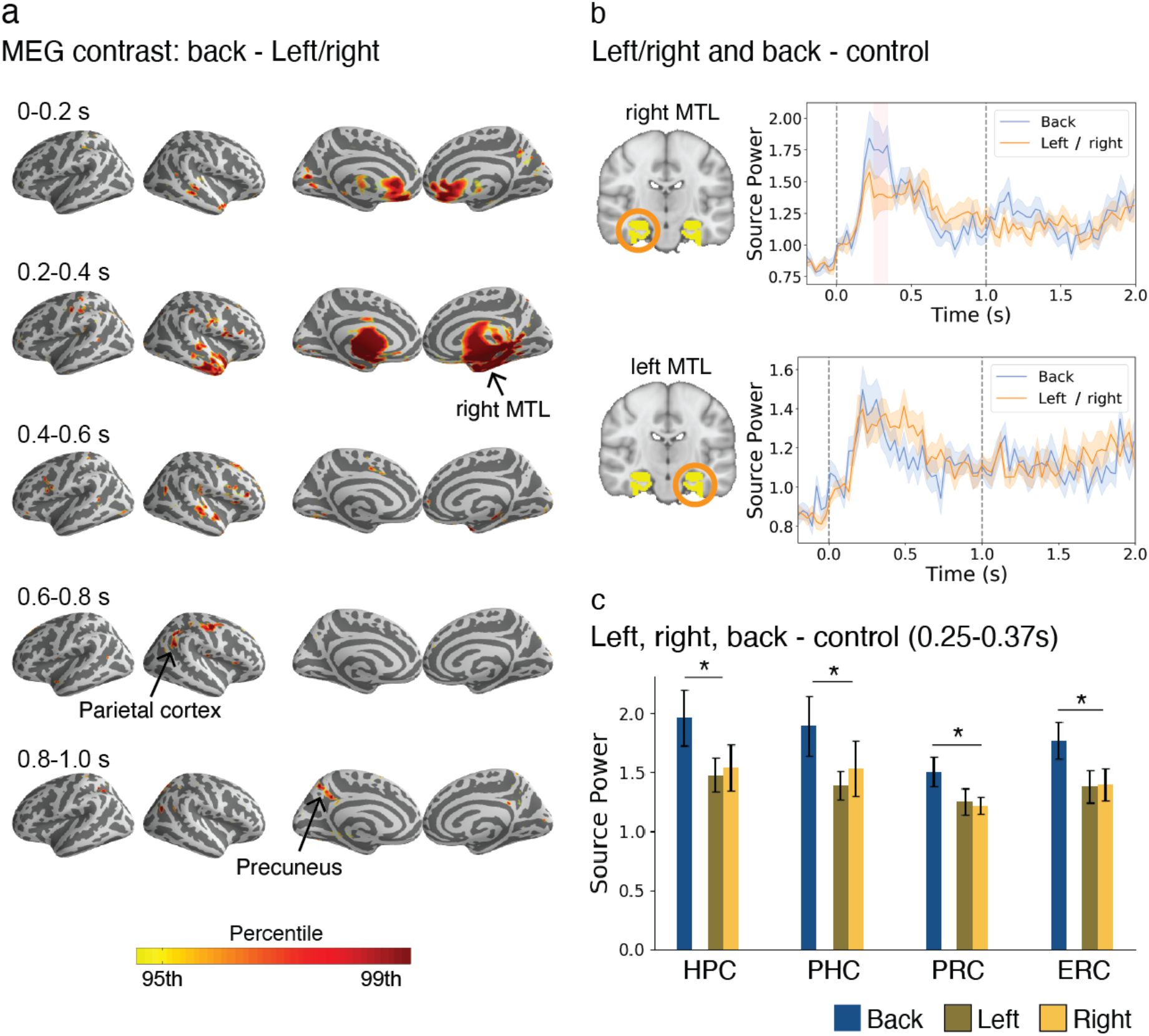
a) MEG contrast for the “back-target” – “left/right-target” condition in source-power for every 0.2 s during the targeting period. The color bar represents the percentile rank of source-power strength. b) Time courses of the source-power in the anatomical ROIs for both hemispheres of the MTL. The shaded area (right MTL, 0.25–0.37 s after the onset of targeting period in the top panel) indicates a significantly higher source-power in the back-target condition than in the left/right-target condition (*P* < 0.05, spatial-temporal cluster correction for multiple comparison, two tailed). c) ROI analysis of the source-power in each of the right MTL subregions for each condition. \**P* < 0.05, *t* (11) = 3.00, 2.98, 3.22, and 3.39 for HPC, PHC, PRC, and ERC, respectively, “back” vs. “left/right,” two-tailed, Bonferroni-corrected for multiple comparisons (n = 4). Error bars represent SEMs.

### MEG functional-connectivity analysis

Next, we examined the connectivity of the parietal cortex with the FEF, SMA, and MTL areas by calculating the phase lag index (PLI) for the MEG data^31–32^. We chose the two time-windows of interest (0.08–0.48 s and 0.56–0.96 s after the onset of targeting period, see method for details) since they could include at least three cycles of alpha-band waves and cover the early and late phases, which were revealed from the MEG contrast analysis (Figs. 5&6). Figure 7 shows the differences in connectivity between the back- and left/right-target conditions. The connectivity patterns differed significantly between the two time windows in the alpha band (8–13 Hz) [*F* (1, 132) = 8.24, *P* = 0.02, repeated-measures two-way ANOVA, Bonferroni-corrected for multiple comparisons, n = 4 for frequency bands], but not in the other bands. During the early time window, the parietal cortex showed a larger connectivity with the right ERC and PRC of the MTL under the back-target condition than under the left/right-target condition, although the difference was statistically marginal [ERC: *t*(11) = 2.16, *P* = 0.06; PRC: *t*(11) = 1.91, *P* = 0.08, uncorrected]. Conversely, we found a larger connectivity of the parietal cortex with the FEF [*t*(11) = 2.61, *P* = 0.02, uncorrected] and SMA [*t*(11) = 1.73, *P* = 0.11, uncorrected] under the left/right-target condition than under the back-target condition during the late time window. These results were consistent with those obtained with functional connectivity analysis using fMRI (Fig. 4). In contrast to the time-dependent alpha-band connectivity of the parietal cortex to the MTL and frontal areas, we did not find a change in the connectivity of the precuneus with any of the ROIs in any frequency band across the two time windows of interest.

**Figure 7.**
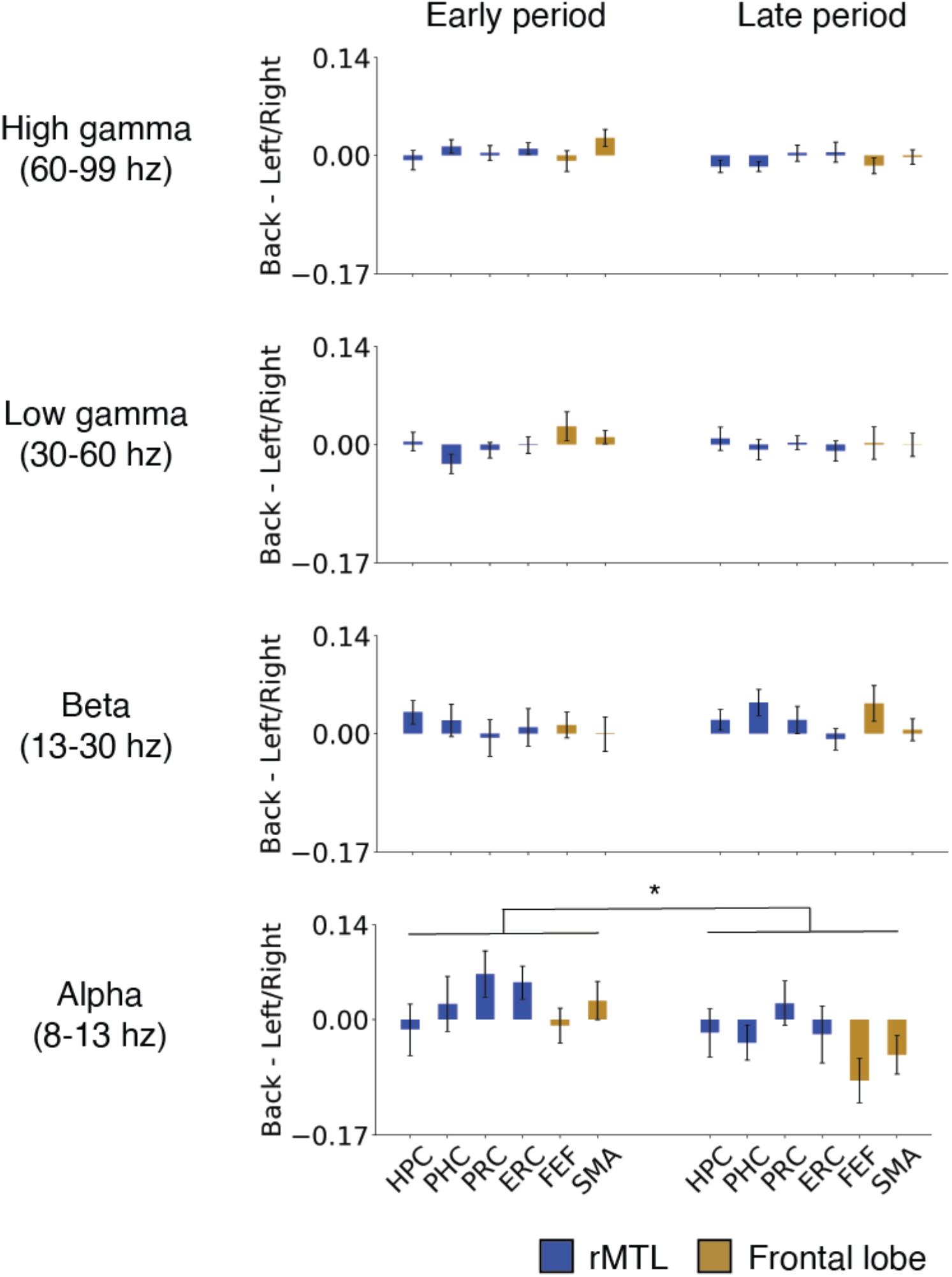
MEG connectivity contrast for the “back-target” – “left/right-target” condition using the parietal cortex as a seed. The connectivity with each of the six anatomical ROIs was estimated for alpha, beta, and gamma frequency bands using PLI during the early phase (0.08–0.48 s) and late phase (0.56–0.96 s). \**P* = 0.02, *F* (2, 132) = 8.24, a main effect of time-windows (early vs. late), repeated-measures two-way ANOVA with brain areas as another main effect, Bonferroni-corrected for multiple comparisons of frequency bands (n = 4). Error bars represent SEMs.

## Discussion

The present combined fMRI and MEG study showed greater involvement of the fronto-parietal network when a participant remembered a target location on the left/right side than on the back of the self-body. Conversely, larger involvement of the MTL-parietal network was revealed when a target object was behind a participant. These results suggest that the parietal cortex represents the external space surrounding the self-body in mind by interacting with the dorsal frontal regions and the MTL.

A previous human fMRI study using multi-voxel pattern analysis (MVPA) reported that the parietal cortex codes the egocentric space both inside and outside the visual field^28^, which is consistent with the findings of our previous MVPA-based study that employed the same spatial-memory task as the present one^29^. However, Schindler and Barteles^28^ did not find the relationship between BOLD signal strength and egocentric space demonstrated by the present study. One reason for this discrepancy might be the different types of memory required for the two studies. Schindler and Barteles^28^ intensively trained participants to learn a fixed spatial arrangement of eight objects for several days before the MRI scans. During the scans, the participants remembered a target object location based on the knowledge that they stored as long-term memory in a different room. In contrast, the present study prompted the participants to encode a trial-specific spatial relationship among the three objects in each trial (see Methods) and remember a target location from their short-term memory in the same 3D virtual environment as they encoded during the scan. These different task conditions may have resulted in different participant retrieval strategies between the two studies, which may have affected space representations in the parietal cortex.

The present study exhibited disproportional spatial representations around the self-body, with a bias toward the left/right-target location in comparison with the back-target location. The preferential processing of the left/right-target location relative to the back by the frontoparietal network is consistent with the results of previous behavioral studies that reported “front facilitation,” i.e., the location of a target in the anterior part of a participant is more efficiently detected than a target located behind them^24,25,27^. On the other hand, we found greater involvement of the MTL-parietal network in representing the back target location than the left/right target location. Interestingly, the present MEG experiment comparing each target-location condition with the control condition showed that the source power increased in the MTL not only for the back-target location but also for the left/right-target location (Figs. 6b). These results suggest that the retrieved target location information is transmitted from the MTL to the parietal cortex regardless of whether the location was inside or outside the visual field. These findings are consistent with the results of previous studies that suggested the involvement of both the MTL and parietal cortex as members of the core brain system in the recollection of episodic memory^33,34^. We hypothesized that the MTL-parietal network was involved in the back-target condition more than the left/right-target condition to generate a mental representation of a retrieved target location because the representation in the left/right-target condition was also supported by the frontoparietal network, which might reduce the demand on the MTL-parietal network in comparison with the back-target condition.

While the fMRI experiment suggested the preferential involvement of only the rhinal cortex, particularly the ERC in the back-target condition, the MEG experiment suggested the involvement of all MTL subareas. This inconsistency might be due to the ill-posed nature of the MEG inverse problem (e.g., the “source leakage”), since a limited number of magnetic-field sensors yielded insufficient activity to discriminate among thousands of source points, particularly for neighboring regions^35–37^. The MEG source power in the other MTL subregions might thus be caused by signal leakage from the ERC. Another possibility might be that the MEG signal reflected synchronized activity at each instantaneous time point and would be more sensitive to transient neuronal operations than fMRI analysis, which is based on the averaged BOLD signal^38^ in each TR (2 s). In either case, the ERC might play a key role in the MTL-parietal network for the representation of a retrieved target location, which is consistent with the results of previous studies that examined the spatial properties of the ERC neurons (e.g., grid cells, head direction cells) in both rodents^22,39–42^ and primates^43–45^. Importantly, the primate ERC reportedly represents the external space according to the gaze position and even in imagined navigation^46^. In contrast to the ERC, other MTL subregions may be involved in the retrieval of the target location^47–49^. Considering the high performance of the present spatial memory task and the fact that post-scan interviews of the participants showed no strategic efforts for retrieval, the retrieval process might be only transient in the present experimental paradigm, which could be detected more efficiently by MEG than by fMRI.

The MEG connectivity analysis using PLI revealed a preferential increase in synchronization at alpha-band frequency (8–13 Hz) for both frontoparietal and MTL-parietal networks for the left/right-target and back-target conditions, respectively. In contrast, we did not observe a differential increase in synchronization in the other frequency bands. These findings were consistent with the results of previous MEG and EEG studies, which suggested functional roles of alpha-band phase synchrony in long-range communications across distant brain regions, including the parietal cortex^8,50–53^. One remaining question concerns the functional significance of the MTL-parietal network in daily life. In the present spatial-memory paradigm, the participants obtained the egocentric target location from their short-term memory, either inside or outside the visual field^29^. Instead, we usually locate a target inside the visual field for a subsequent action by looking at it^54,55^, while we occasionally retrieve a target behind us from short-term memory. Therefore, the visual modality of information on the back is always encoded in the past, and the preceding spatiotemporal information is retained in the MTL^56,57^. Collectively, the parietal cortex may represent the whole space around the self-body by coordinating the MTL-parietal network and the frontoparietal network, which may equip us with the mental representation of the present external world interacting with our past (retrieval) and future (action planning).

## Materials and Methods

### Participants

Nineteen and 12 right-handed university students with normal or corrected-to-normal vision were recruited from Peking University for fMRI and MEG experiments, respectively (fMRI: 12 women, 7 men; MEG: 4 women, 8 men). The average age of the participants recruited for the fMRI and MEG experiments was 24.9 years (range: 18–30 years) and 22.5 years (range: 19–25 years), respectively. None of the participants had a history of psychiatric or neurological disorders; all of them provided written informed consent prior to the start of the experiment, which was approved by the Research Ethics Committee of Peking University.

### Experimental design

#### Experimental design

The details of the study design have been described previously^29^. A 3D virtual environment was programmed using the Unity platform (Unity Technologies, San Francisco). Three animated 3D human characters (Mixamo, San Francisco, https://www.mixamo.com) were placed on three out of four locations pseudo-randomly across trials (Fig. 1a). Participants performed the task using a first-person perspective with a 90° field of view (aspect ratio = 4:3) and had never seen a top-down view of the virtual environment. Experimental stimuli were presented through an LCD projector with a resolution of 1024 × 768 pixels.

The spatial memory task included 144 and 72 trials in the MRI and MEG experiments, respectively. In each trial, participants walked from one of the four starting locations toward the human characters and stopped on the wood plate in the center of the environment. After the walking period, participants experienced a “facing period” and a “targeting period” sequentially. In the facing period, one of the human characters was presented as a facing-character in the center of the display with the environmental background for 2.0 s (MRI) or 1.0 s (MEG), with the other two characters being invisible. In the targeting period, a photo of one of the remaining characters (named as the “targeting-character”) was presented as a target on a scrambled background for 2.0 s (MRI) or 1.0 s (MEG). Each of the three periods was followed by a 2.0-s (MRI) or 1.0-s (MEG) delay (noise screen). At the end of each trial, participants indicated the direction of the target relative to their self-body by pressing a button when a cue was presented.

We added head-nodding detection (HND) trials (16 trials for MRI and 36 trials for MEG) to the spatial memory task. In the HND trials, a photo of one of the human characters was presented after the walking period, and the participants were then asked to indicate whether the human character nodded its head during the walking period. Each human character nodded its head at a probability of 20.6% at a random time point between the start and end of walking. Because the trial types were indistinguishable during the walking period, the participants were required to pay attention to the head-nodding of the human characters during the walking period, which would reduce the possibility of voluntary memorization of the spatial relationship of the three objects. Post-scanning interviews showed that none of the participants made efforts to memorize the spatial relationship^29^.

#### MEG control conditions

Two control conditions (36 trials for each) were added to the spatial memory task in the MEG experiment. In the control conditions, after the walking period, a white cross was presented instead of a human character during both the facing and targeting periods or during the targeting period (Fig. 4c). The participants were instructed to rest with their eyes open and fixate on the white cross. After the targeting period, the participants pressed a button corresponding to the number presented on the screen during the response period. The number was determined randomly from one to four in each trial of the control conditions.

### fMRI acquisition and analysis

#### MRI scanning parameters

BOLD MRI images were acquired using a 3T Siemens Prisma scanner equipped with a 20-channel receiver head coil. Functional data were acquired with a multi-band echo planar imaging sequence (TR: 2000 ms; TE: 30 ms; matrix size: 112 × 112 × 62; flip angle: 90°; gap: 0.3 mm; resolution: 2 × 2 × 2.3 mm^3^; number of slices: 62; slice thickness: 2 mm; gap between slices: 0.3 mm; slice orientation: transversal). The signals of the original voxels (i.e., 2 × 2 × 2 mm^3^) were assigned to the corresponding voxels without a gap (2 × 2 × 2.3 mm^3^) to construct participants’ native space images. Four experimental sessions were conducted with 478, 476, 473, and 475 of TRs on average. A high-resolution T1-weighted three-dimensional anatomical data set was collected to facilitate registration (MPRAGE: TR: 2530 ms; TE: 2.98 ms; matrix size: 448 × 512 × 192; flip angle: 7°; resolution: 0.5 × 0.5 × 1 mm^3^; number of slices: 192; slice thickness: 1 mm; slice orientation: sagittal).

#### fMRI preprocessing

BOLD images of each scanning session were preprocessed independently using FSL FEAT (FMRIB’s Software Library, version 6.00, https://fsl.fmrib.ox.ac.uk/fsl/fslwiki)^58,59^. For each session, the first three functional volumes were discarded to allow for T1 equilibration, and the remaining functional volumes were slice-time corrected, realigned to the first image, high-pass filtered at 100 s, and smoothed using a 5-mm FWHM Gaussian filter.

#### Univariate analysis

The 4-s BOLD signals of the targeting period were modeled using univariate general linear models (GLM) with the three egocentric directions (left, right, and back) included as regressors in each experimental session (40 trials) for each participant. The BOLD signals in the other task periods were modeled by additional regressors, including 12 visual patterns for the 8-s walking period (three spatial arrangements of human characters × four walking directions), the four types of body rotation for the 4-s facing period (135° and 45° in the clockwise and anti-clockwise directions), the three buttons that participants pressed during their response for the response period, the time period for the HND trials, and the six motion parameters. A canonical hemodynamic response function was used for each task period in GLM. This procedure generated a parameter map for each of the three egocentric directions (left, right, and back). The parameter maps were averaged across four scanning sessions for each egocentric direction and registered to a T1-weighted standard image (MNI152) using FSL FLIRT^60,61^ before group-level statistical analyses. The voxel size was converted into a 2 × 2 × 2 mm^3^ resolution during the registration process.

#### Connectivity analysis

To conduct whole-brain analyses examining the functional connectivity of the parietal cortex and precuneus, we first removed several sources of spurious variance along with their temporal derivatives from the preprocessed functional data by performing a GLM, which included (1) the mean time courses of the BOLD signal across voxels within the lateral ventricles; (2) white matter; (3) whole brain; and (4) the six motion parameters as regressors. The residual signals were bandpass-filtered, leaving signals within the frequency range of 0.01–0.1 Hz and shifted by two TR intervals (4 s)^62–64^. The signals in the participant’s native space were registered to a T1-weighted standard image (MNI152) using FSL FLIRT^60,61^. Then, we extracted the TRs during the 4-s targeting period from each trial and concatenated them with those in the next trial for each of the three egocentric directions and each session. The BOLD signals in each voxel were z-transformed along the time course for each direction. Next, we averaged the z-transformed BOLD signals over the anatomical masks of the brain areas (i.e., the seeds of the parietal cortex and precuneus) on the basis of the automated anatomical labeling (AAL) template^65^, which generated a regional time course across trials for each egocentric direction in a session^66^. The regional time course of each seed was then correlated with the time course of each voxel in the whole brain, which yielded a whole-brain correlation map for the left-, right-, and back-target conditions separately. The correlation maps were averaged across four scanning sessions (and left-and right-target conditions) before submitting them to a group-level statistical analysis.

### MEG acquisition and analysis

#### MEG scanning parameters

Neuromagnetic signals were recorded with a 275-channel whole-head axial gradiometer DSQ-3500 MEG system (CTF MEG, Canada) at a sampling rate of 1200 Hz. A third-order synthetic gradiometer and linear drift corrections were applied to remove far-field noise. To measure the head position within the MEG helmet, three head-position indicator (HPI) coils were attached to the nasion and two preauricular points of each participant to coregister their head position with the sensor coordinate system. During scanning, customized chin-rest equipment compatible with MEG was prepared to ensure that head movements did not exceed 2 mm. After MEG recording, each participant underwent anatomical MRI scans on a 3T Siemens Prisma scanner (voxel size: 1 mm^3^; flip angle: 9°; TE: 1.97 ms; TR: 2,300 ms; field of view: 256 × 256 × 176 mm^3^), and three MRI markers were attached to the locations of the HPI coils to align each participant’s anatomical image to the MEG sensor positions.

#### MEG preprocessing

Raw signals from the MEG experiment were preprocessed and analyzed using the MNE Python toolbox (v0.19; available at: https://mne.tools/stable/index.html)^67,68^. The time courses of the MEG signal were bandpass-filtered between 1 and 100 Hz offline. Artifacts (e.g., eye movements, eye blinks, and cardiac movements) were removed by performing independent component analysis (ICA). Then, the MEG signals were visually inspected and downsampled to 200 Hz to increase the processing speed in later analyses^69^. *Sensor space analysis:* The baseline MEG signal for each trial was calculated by averaging the signals during the 0.2-s period preceding the onset of the targeting period, and the baseline was subtracted from the signal at each time point during the targeting period in that trial. Then, the time series of the MEG signal was averaged across trials for each target and control condition. Contrast analyses were performed to compare the difference between the left/right-target and back-target conditions, and to test the effects of the recollection in each of the three target conditions relative to the control conditions. These analyses were performed on each sensor, and a sensor space was generated for each session. The sensor spaces were averaged across sessions before submitting them to the statistical test for group-level statistical analysis.

#### Source space analysis

To reconstruct the spatial-temporal activity from the sensor space to the anatomical space, the forward model was created using a single-compartment (inner skull) boundary-element method on the basis of each participant’s anatomical image, and the spatial-temporal activity was then inversely modeled using the dynamic statistical parameter map at each source point and time^30^. The source space was estimated using a subsampling strategy, which involved subdividing a polygon (oct6) using the spherical coordinate system provided by FreeSurfer, producing 4098 source points per hemisphere with an average source spacing of 4.9 mm (assuming a surface area of 1000 cm^2^/hemisphere)^67,70^. The source space of each participant was morphed to an average surface. The percentile ranks of source-power strength from the top 5% to 1% was calculated for each egocentric location in comparison with the control condition or the contrast between the left/right and back conditions. For ROI analysis, we manually delineated each of the medial temporal lobe (MTL) subareas (hippocampus [HPC], parahippocampal cortex [PHC], perirhinal cortex [PRC], and entorhinal cortex [ERC]) on the participant’s native space using established protocols^71–74^ as well as the delineating software ITK-SNAP (www.itksnap.org). The mean source power within the MTL subareas was calculated by averaging the source power within each mask for each egocentric location before submitting it to a group-level statistical analysis.

#### Connectivity analysis

To assess the connectivity of the parietal cortex and precuneus with the frontal lobe (FEF/supplementary motor area [SMA]) and MTL subareas in each temporal period revealed by the MEG contrast analysis (i.e., “early” [0.25–0.37 s after the onset of targeting period] for the back-target condition and “late” [0.67–0.85 s] for the left/right-target condition), we examined the phase synchronization of the MEG time series for the following four frequency bands: alpha (8–13 Hz), beta (13–30 Hz), low-gamma (30–60 Hz), and high-gamma (60–99 Hz) bands. For this purpose, we first calculated the powers of the MEG signal in a time window of 400 ms centering iteratively at each of the time points within the two time periods. The 400-ms time window was applied in this analysis to ensure that at least three cycles of the source time series could be covered in the alpha band^75^. The powers at each time point were averaged across trials including all target locations and a control condition for each participant, and the grand-mean of power across participants was calculated. We selected the centering time point of the time window that had the maximum grand-mean power for each time period. This procedure yielded two time-series (the time window of 0.08–0.48 s corresponding to the centering point of 0.28 s and the time window of 0.56–0.96 s corresponding to the centering point of 0.76 s for the “early” and “late” periods, respectively). Phase synchronization was tested among the eight ROIs (parietal cortex, precuneus, FEF, SMA, and right MTL subareas) for each time series, condition (left, right, or back), and frequency band using PLI^31,32^ and the built-in function of the MNE Python toolbox^67,68^. In each frequency band, we calculated the difference in the connectivity of each time series for each ROI between the back- and left/right-target conditions before group-level statistical analysis.

### Statistical analysis

For the MRI univariate and connectivity analyses, an initial threshold of *P* < 0.01 was applied to each voxel, and the reliability of clusters was tested using a nonparametric statistical inference that did not make assumptions about the distribution of the data^22,66,67^. The test was conducted with the FSL randomize package (version v2.9, http://fsl.fmrib.ox.ac.uk/fsl/fslwiki/Randomise) with 5000 random sign-flips, and clusters with a size higher than 95% of the maximal supra-threshold clusters in the permutation distribution were then reported. Data obtained by ROI analysis of the MRI BOLD signal, MEG source power, and MEG connectivity analysis were tested using either a *t*-test with Bonferroni correction or repeated-measures two-way ANOVA. The MEG sensor space analysis used either a spatial-temporal cluster permutation test or a spatial-cluster permutation test. All statistical tests were two-sided unless otherwise noted, and significance was determined when the corrected *P* value was less than 0.05.

## Data and code availability

The datasets and code supporting the current study are available from the corresponding author (Yuji Naya) upon reasonable request.

## Acknowledgments

This work was supported by the National Natural Science Foundation of China Grant 31421003 (to Y.N.) and by the Fundamental Research Funds for the Central Universities, PKU (7100602954 to Y.N.). We thank the National Center for Protein Sciences at Peking University for assistance with MRI data collection, and the State Key Laboratory of Brain and Cognitive Science at the Chinese Academy of Sciences for assistance with MEG scanning, which was supported in part by grants from the Ministry of Science and Technology of China (2019YFA0707103), National Nature Science Foundation of China (31730039), and Strategic Priority Research Program of the Chinese Academy of Science (XDB32010300). Computational work was supported by resources provided by the High-performance Computing Platform of Peking University.

